# *Lactobacillus acidophilus* Attenuates Polyethylene Glycol-Induced Susceptibility to *Citrobacter rodentium* Infection via Microbiota Modulation

**DOI:** 10.1101/2025.02.11.637759

**Authors:** Guan-jun Kou, Jing Shen, Guang-chao Li, Shan-shan Fu, Li-xiang Li, Xiu-li Zuo, Yan-qing Li

## Abstract

**Objective:** Polyethylene glycol (PEG), a globally prevalent laxative, is extensively utilized for bowel preparation prior to colonoscopy or abdominal surgery owing to its exceptional safety profile. Nevertheless, emerging evidence suggests that PEG administration may induce significant alterations in colonic microbiota composition, with dysbiotic effects persisting for over two weeks. The potential implications of such microbiota perturbations on the colonization and invasion of opportunistic pathogens remain underexplored.

**Methods and Analysis:** To investigate this, we employed a murine model infected with *Citrobacter rodentium* (*C.R.*), a surrogate for E*nteropathogenic Escherichia coli* (EPEC) and *Enterohemorrhagic Escherichia coli* (*EHEC*), following PEG-induced bowel preparation.

**Results:** The severity of infectious enteritis was assessed, and the prophylactic efficacy of *Lactobacillus acidophilu*s (*LAC*) supplementation was evaluated. Our findings indicate that PEG administration significantly elevated *C.R.* load, enhanced virulence gene expression, and exacerbated intestinal inflammation post-infection, with the infection window extending approximately 14 days. Through fecal metagenomic analysis and co-housing experiments, we demonstrated that PEG-associated enteritis is critically mediated by gut microbiota dysbiosis. Furthermore, supplementation with *LAC* was shown to mitigate susceptibility to *C.R*. infection.

**Conclusions:** These results suggest that PEG increases intestinal susceptibility to infection in a microbiota-dependent manner, highlighting the therapeutic potential of *LAC* in restoring microbial homeostasis.

## Introduction

Polyethylene glycol (PEG) is extensively utilized for comprehensive bowel preparation prior to colonoscopy or abdominal surgery, owing to its exceptional safety profile [1]. Previous investigations have demonstrated that PEG consumption significantly alters the composition of the intestinal microbiota [2]. Specifically, PEG administration modifies the α- and β -diversity of the intestinal microbiota within 6 days, increases the prevalence of mucus-associated bacteria (e.g., *Akkermansia*, *Bacteroides*), and reduces the abundance of *Firmicutes* after 14 days [3]. However, another study indicated that the abundance of gut microbiota in volunteers following PEG-induced bowel cleansing was 34.7-fold lower than baseline levels, with recovery to normal levels occurring approximately two weeks later [4]. These findings underscore the need for further research into the effects of PEG on intestinal microbiota and the rate of bacterial recovery.

A previously published meta-analysis revealed that laxative-induced alterations in the intestinal flora are comparable to those induced by proton-pump inhibitors (PPIs) [5]. The use of PPIs has been positively correlated with an increased risk of enterovirus infection and a 1.8-fold elevated risk of *Clostridium difficile* infection [6]. Consequently, we hypothesize that bowel cleansing with PEG electrolyte solution may similarly enhance susceptibility to enteric infections.

*Enteropathogenic Escherichia coli* (*EPEC*) and *Enterohemorrhagic Escherichia coli* (*EHEC*) are primary pathogenic bacteria responsible for infectious enteritis in humans [7]. *Citrobacter rodentium* (*C.R.*) is a rodent-specific enteric pathogen capable of adhering to the colonic mucosa and forming attachment and effacing (A/E) lesions, including colonic crypt hyperplasia, which closely resembles the pathology induced by *EPEC* and *EHEC* [8]. As such, *C.R.* is frequently employed as a model organism to study the mechanisms of intestinal infection in murine systems [9–13].

*Lactobacillus acidophilus* (*LAC*) is among the most widely used probiotics. It has been demonstrated to not only ameliorate gut dysbiosis, as evidenced by reduced *Firmicutes*-to-*Bacteroidetes* ratios and decreased levels of endotoxin-bearing Gram-negative bacteria, but also to maintain intestinal barrier integrity and reduce inflammation [14]. These properties suggest that *LAC* may play a protective role in mitigating the adverse effects of PEG-induced microbiota alterations and subsequent enteric infections. Further studies are warranted to elucidate the potential therapeutic benefits of *LAC* in this context.

## Materials and Methods

### Mice

Wild-type C57BL/6 male mice (6-7 weeks old) were obtained from Jicui Biological Technology Co., Ltd. (Nanjing, China). The animals were housed under specific pathogen-free (SPF) conditions, maintained at a controlled temperature of 23 ± 1°C and relative humidity of 45 ± 5%. Following a one-week acclimatization period, the mice were utilized for experimental procedures. All animal experiments were conducted in accordance with protocols approved by the Institutional Animal Care and Use Committee of Qilu Hospital, Shandong University (Approval No. Dull-2021-058).

### PEG administration

Intestinal cleansing was performed using 2.1 mL of PEG electrolyte solution (Heshuang, Wanhe Pharmaceutical, Shenzhen, China; containing PEG 4000 as active ingredient). Control groups received either 2.1 mL of sterile distilled water. Administration was performed via oral gavage with 300 μL aliquots of the respective solutions at 30-minute intervals, followed by a 4-hour fasting period.

### *C.R.* infection model

C.R. strain DBS 100 was cultured overnight in Luria-Bertani (LB) broth at 37°C under aerobic conditions. Bacterial cells were harvested by centrifugation at 4000 × g for 10 minutes at 4°C. The pellet was washed twice with PBS through sequential centrifugation (4000 × g, 3 minutes) and resuspended in sterile PBS. Mice were infected via oral gavage with 200 μL PBS containing 1 × 10^9 colony-forming units (CFUs) of C.R.

For bacterial quantification, fecal samples were collected at designated time points post-infection and homogenized in PBS. Tissue samples, including colon, mesenteric lymph nodes (mLNs), spleen, and hepatic parenchyma, were aseptically collected and weighed.

Tissue homogenates and fecal suspensions (30-100 μL) were plated on MacConkey agar and incubated at 37°C for 14-16 hours. *C.R.* colonies were enumerated and expressed as CFUs per gram of tissue or fecal material.

### *LAC* administration

Following PEG administration, mice were orally gavaged daily for 3 consecutive days with either 1.86 × 10^6^ CFU of *LAC* (Hangzhou Grand Pharmaceutical Co., Ltd., China) suspended in 200 μL PBS or 200 μL PBS alone (designated as the PBS control group). On Day 3 post-treatment, all animals underwent standardized *C.R.* infection protocol as previously described. Experimental groups comprised: Control group (n = 5) and PEG group (n = 5).

### DNA or RNA extraction and quantitative real-time PCR

For microbial genomic profiling, freshly voided fecal specimens were aseptically collected in RNase/DNase-free microcentrifuge tubes and flash-frozen in liquid nitrogen prior to storage at -80°C. Total bacterial genomic DNA was purified using the Stool Genomic Plus DNA extraction kit (Juhemei Biotechnology Co., Ltd., China) following manufacturer’s specifications.

Concurrently, distal colonic mucosal biopsies were homogenized in RNA isolation reagent (Vazyme, R401-01), with subsequent RNA integrity verification via agarose gel electrophoresis. RNA quantification was performed using a NanoDrop™ 2000 spectrophotometer (Thermo Fisher Scientific). First-strand cDNA synthesis was carried out using 1 μg total RNA with HiScript III RT SuperMix for qPCR (+gDNA wiper) (Vazyme, R323-01), incorporating genomic DNA elimination steps.

Quantitative PCR analyses were conducted on a StepOnePlus™ Real-Time PCR System (Thermo Fisher; 4376600) utilizing the following reaction parameters: 95°C for 60 s, followed by 40 cycles of 95°C for 15 s, 60°C for 15 s and 72°C for 45 s. Reaction mixtures (20 μL total volume) contained: 10 μL 2× AceQ® Universal SYBR qPCR Master Mix (Vazyme, Q511-02), 1 μL cDNA template, 0.8 μL each of gene-specific forward/reverse primers (Table 1), and 7.4 μL DEPC-treated water. The primers used for qPCR are listed in Table 1. RrsA (16s RNA) was used as the internal standard in order to normalize target gene expression. All reactions were performed in technical triplicates, with relative quantification determined by the comparative 2^-△△Ct^ method.

**Table 1.**
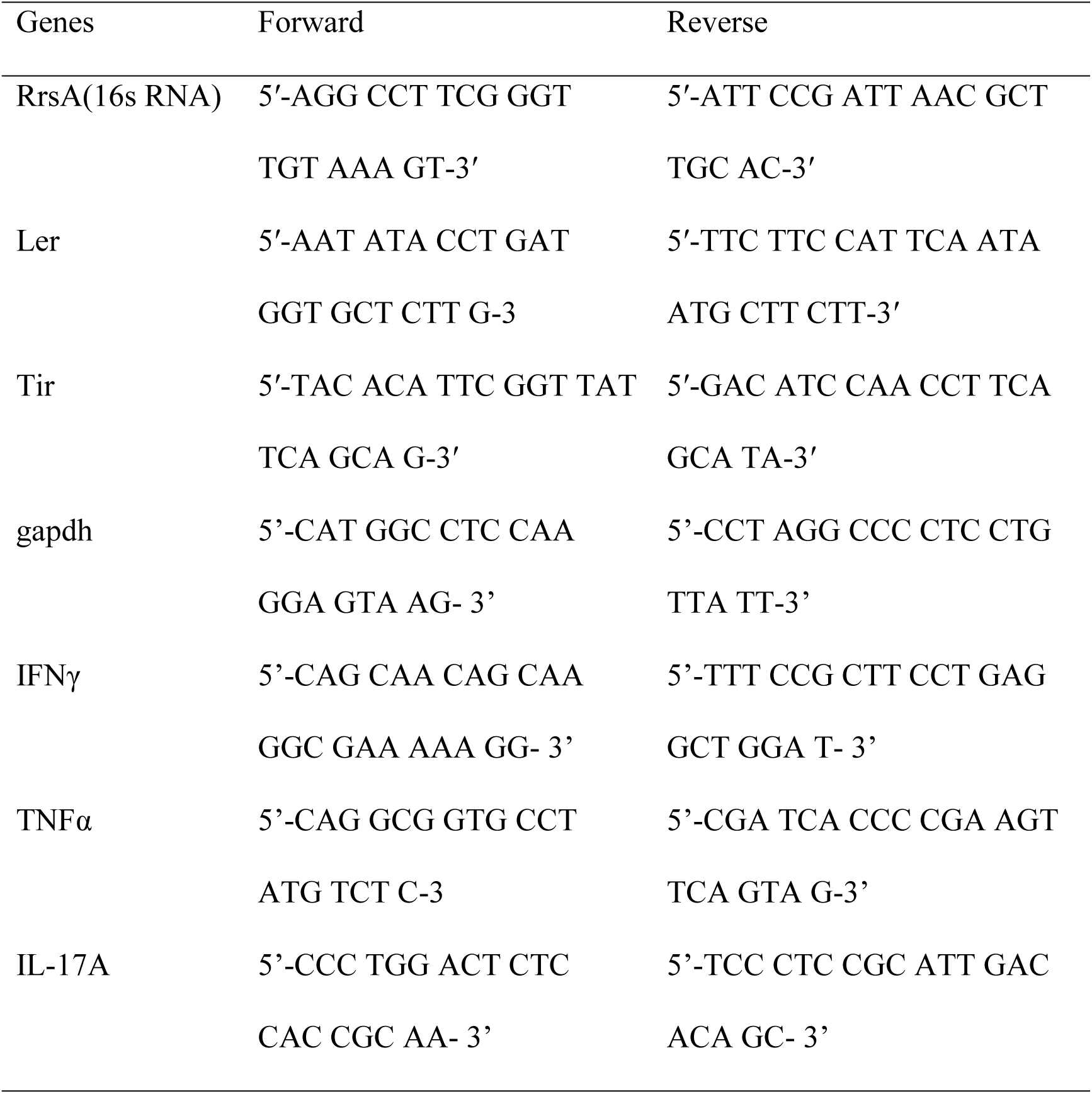
List of primers.

### Hematoxylin and Eosin (H&E) Staining

Approximately 1 cm of distal colon tissue was fixed in 10% neutral buffered formalin at room temperature for 24 hours. Following fixation, tissues underwent a standardized dehydration series, paraffin embedding, and sectioning at 4-μm thickness. Sections were stained with hematoxylin and eosin (H&E) using established protocols. Histopathological scoring was performed in a blinded manner by two independent observers, adhering to the criteria outlined in a previously validated scoring system [15].

### Quantification of goblet cells

To assess the effect of PEG on goblet cell dynamics, mice were euthanized 12 hours post-PEG bowel preparation. Distal colonic tissues were fixed in Carnoy’s fixative (60% methanol, 30% glacial acetic acid, and 10% chloroform) at 4°C for 24 hours, followed by three washes in 70% ethanol. Paraffin-embedded tissues were sectioned at 5-μm thickness and stained with Periodic Acid-Schiff’s stain to visualize mucin-containing goblet cells.

Goblet cells were quantified by analyzing approximately 25 longitudinally oriented crypts per animal using light microscopy (×400 magnification). Data represent biological replicates from 5–7 animals per experimental group.

### Metagenomic Analysis of the Microbiome

Fecal pellets (n = 3 per mouse) were collected aseptically and stored at −80°C until DNA extraction. 16S rRNA sequencing analyses were conducted on the Majorbio Cloud Platform (Shanghai Majorbio Bio-Pharm Technology Co., Ltd.). Microbial genomic DNA was isolated using the E.Z.N.A.® Soil DNA Kit (Omega Bio-Tek, Norcross, GA, USA). The hypervariable V3–V4 regions of the 16S rRNA gene were amplified using universal primers 338F (5′-ACTCCTACGGGAGGCAGCAG-3′) and 806R (5′-GGACTACHVGGGTWTCTAAT-3′) under PCR. Amplicons were sequenced on the Illumina MiSeq platform (2 × 300 bp paired-end; Illumina, San Diego, CA, USA). Raw sequences were processed using UPARSE (v7.1) with a 97% similarity threshold for operational taxonomic unit (OTU) clustering. Taxonomic classification was performed against the Greengenes database.

### Quantification and Statistical Analysis

Data are presented as mean ± standard deviation (SD). Statistical analyses were conducted using two-tailed unpaired t-tests implemented in GraphPad Prism 8 (GraphPad Software, San Diego, CA, USA). The threshold for statistical significance between experimental groups was defined as follows: *p < 0.05, **p < 0.01, and ***p < 0.001.

## Results

### PEG electrolyte solution worsens *C.R.* infection

To investigate the impact of PEG electrolyte solution on *C.R.* infection, wild-type C57BL/6J mice were orally inoculated with *C.R.* following 24-hour pretreatment with PEG. Quantitative analysis revealed that PEG-administered mice maintained significantly elevated fecal *C.R.* burdens (>10⁹ CFU/g) from 3-9 days post-infection (d.p.i.), peaking at day 7 (Figure 1A). In contrast, control animals exhibited approximately 10⁷ CFU/g fecal bacterial loads with similar peak timing (Figure 1B).

Given the established association between intestinal barrier dysfunction and systemic bacterial translocation during enteric infections, we performed comprehensive analysis of bacterial colonization patterns across multiple organ systems. PEG-treated mice exhibited significantly elevated *C.R.* loads in colonic tissues (p<0.01), hepatic parenchyma (p<0.001), splenic tissue (p<0.01), and mLNs (p<0.05) compared to untreated controls (Figure 1C).

**Figure 1.**
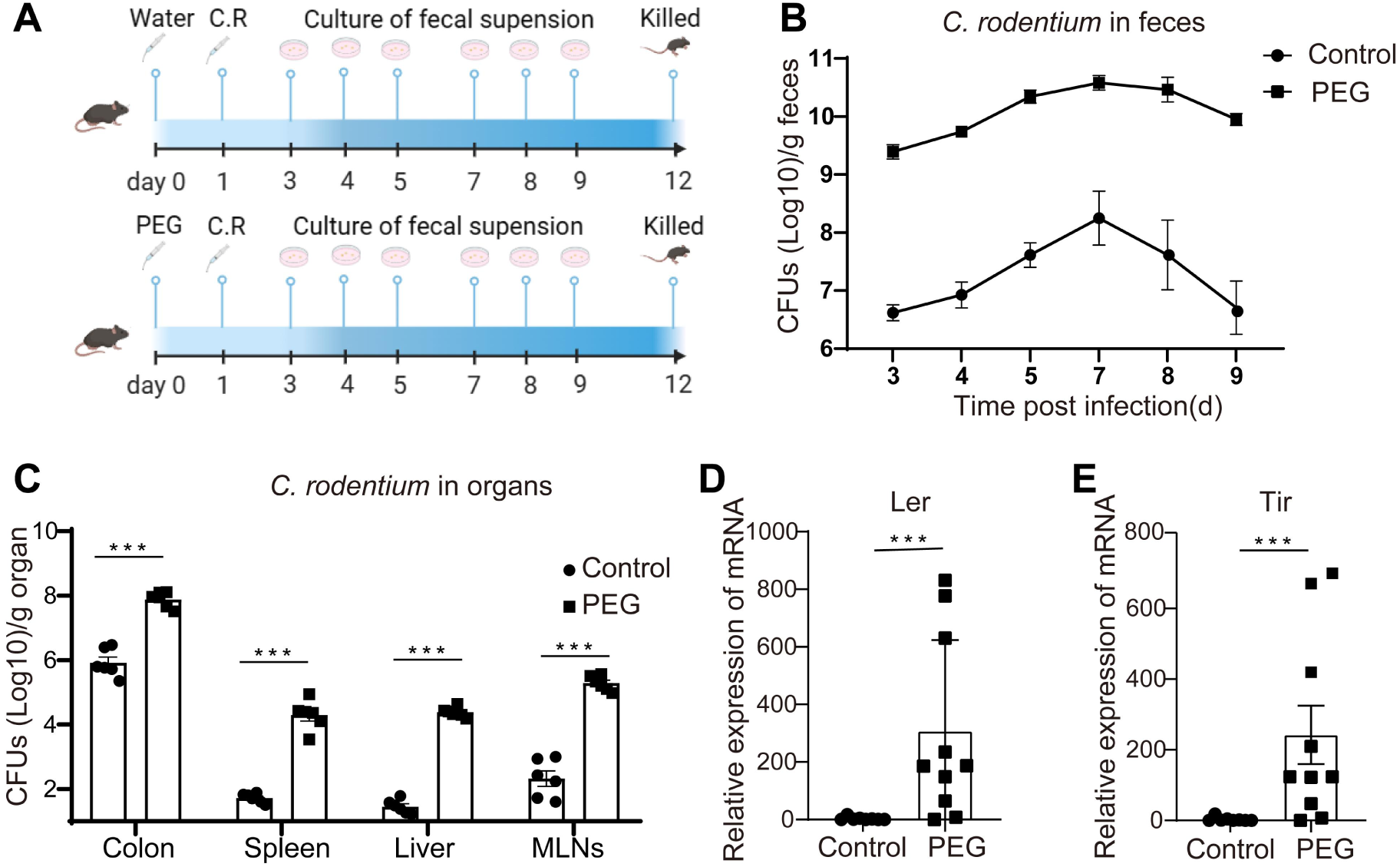
Polyethylene glycol (PEG) increase the *Citrobacter rodentium (C.R.)* load and virulence. (A) Mice of the two groups were gavaged with water or PEG 1 day prior to the infection of *C.R.*. Fecal pallets were collected at day 3, 5, 7, 8, 9. (B) *C.R.* load in the feces over the indicated time. (C) *C.R.* load of the intestines tissue. (D) Systemic dissemination of *C.R.*. (E-F) The mRNA expression of the *Ler* and *Tir* in the fecal.Data are representative of at least two independent experiments. Data are shown as mean ± SD. \**p* < 0.05; \*\**p* < 0.01; \*\*\**p* < 0.001.

To investigate potential PEG-mediated enhancement of *C.R.* pathogenicity, we quantified transcriptional expression of key virulence regulators *Ler* and *Tir* in fecal samples at 5 d.p.i. using qRT-PCR. Notably, PEG administration induced significant upregulation of both ler (91.34-fold increase, p<0.001) and tir (70.87-fold increase, p<0.001) gene expression relative to control animals (Figure 1D).

Histopathological assessment at 12 d.p.i. revealed pronounced colonic shortening in PEG-treated mice compared to controls (8.09 ± 0.33 vs 7.54 ± 0.27, p<0.01; Figure 2A-B). Microscopic evaluation demonstrated exacerbated inflammatory infiltrates and severe crypt architectural disruption in the distal colonic mucosa of PEG-exposed animals, with significantly elevated histopathological scores (6.17 ± 1.17 vs 2.00 ± 0.58, p<0.001; Figure 2C-D).

**Figure 2.**
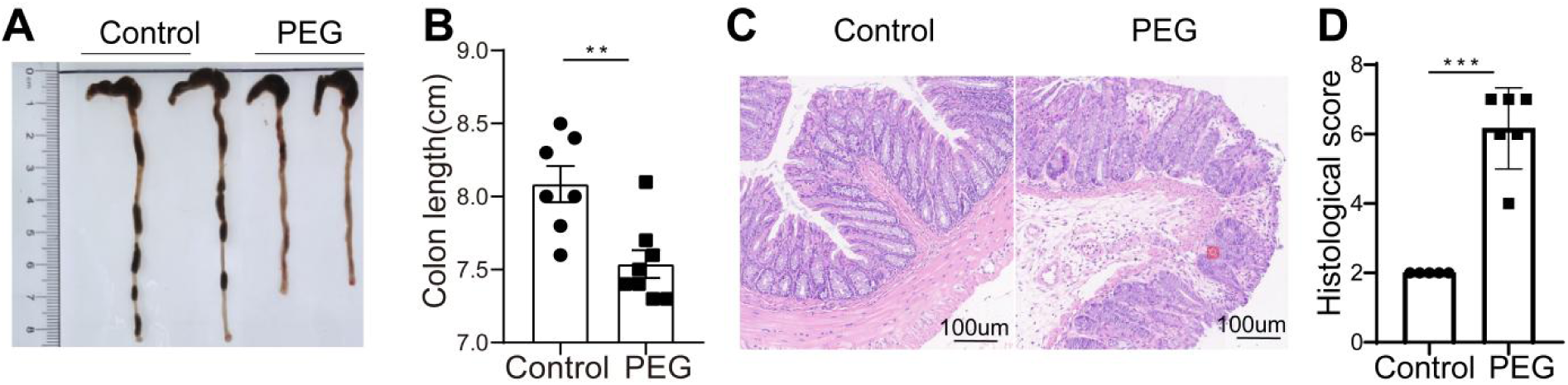
PEG aggravates *C.R.* induced colitis. (A-B) Colon length at day 12 post *C.R.* infection. (C-D) Representative H&E staining (×20) and pathological score of the inflamed colon epithelium. (E-F) Immunoblot for ZO-1 and caludin in colonic tissue and the densitometric analysis of ZO-1 and caludin immunoblot of control and PEG-Streated mice. Data are representative of at least two independent experiments. Data are shown as mean ± SD. \**p* < 0.05; \*\**p* < 0.01; \*\*\**p* < 0.001.

### PEG establishes a 14-day window of infection susceptibility

To temporally characterize PEG-induced intestinal vulnerability, experimental *C.R.* challenges were conducted at 7 or 14 days following PEG-mediated bowel preparation. Bacterial shedding kinetics were assessed through fecal CFU counts at 7 d.p.i., while disease severity parameters (colon morphology, histopathology, and inflammatory markers) were evaluated at 12 d.p.i. (Figure 3A).

**Figure 3.**
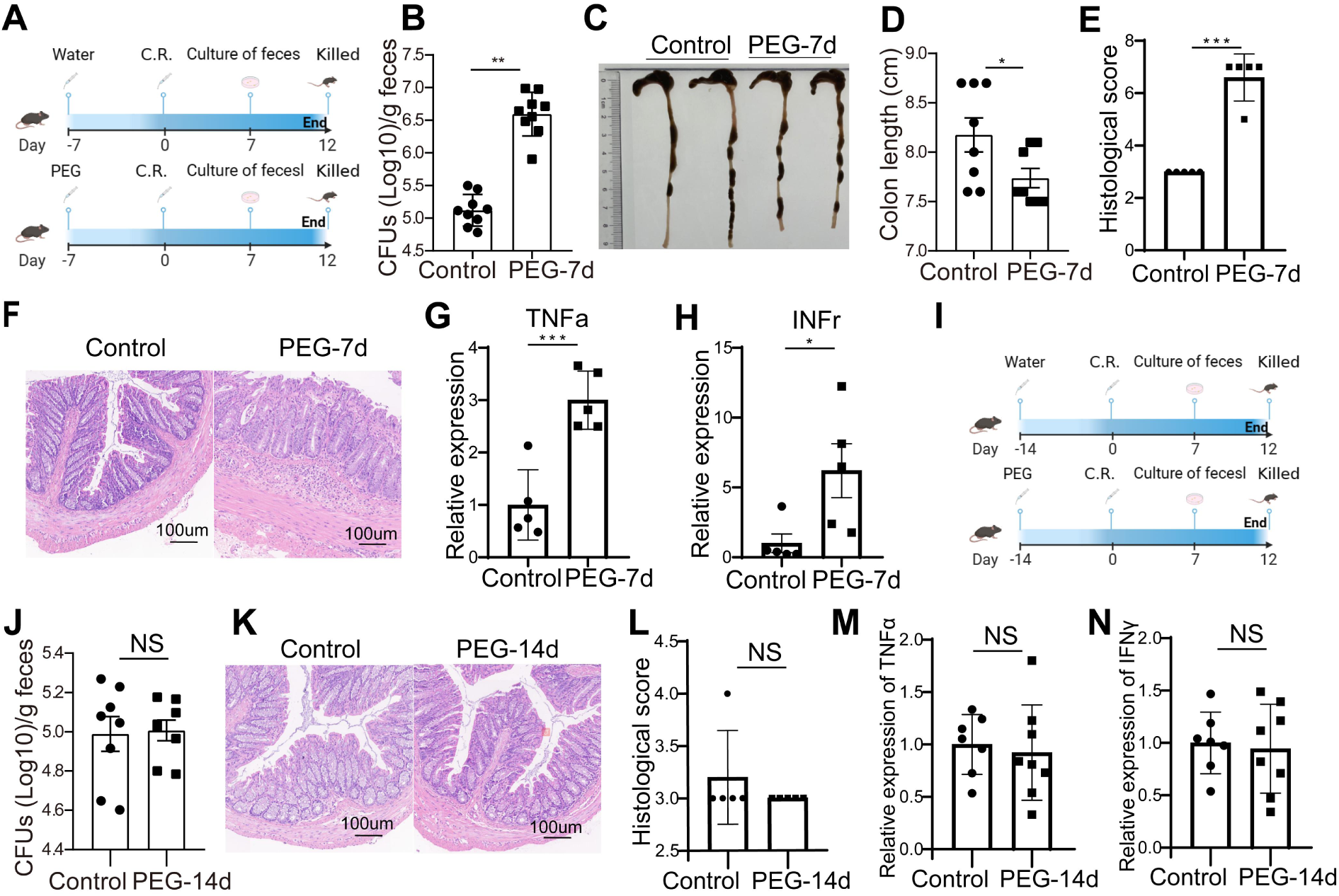
PEG results in a infection window of 14 days. (A) Mice were gavaged with PEG 7-day before infection with C.R. (B) C.R. load of the feces. (C-D) Colon length. (E-F) Representative H&E staining (×20) and pathological score of the inflamed colon epithelium. (G-H) The mRNA expression of the TNFα and IFNγ of the inflamed colon. (I) Mice were gavaged with PEG 14-day before infection with C.R.. (J)C.R. load of the feces. (K-L) Representative H&E staining (×20) and pathological score of the inflamed colon epithelium. (M-N) The mRNA expression of the TNFα and IFNγ of the inflamed colon.Data are representative of at least two independent experiments. Data are shown as mean ± SD. \**p* < 0.05; \*\**p* < 0.01; \*\*\**p* < 0.001. ns, no significance.

Mice infected at 7 days post-PEG treatment displayed enhanced infection severity, manifesting as significantly increased fecal bacterial loads (p<0.001; Figure 3B), marked colonic shortening (p<0.01; Figure 3C-D), and elevated histopathological scores (p<0.001; Figure 3E-F). Quantitative PCR analysis revealed substantial upregulation of proinflammatory cytokine expression in the PEG-7d group, with tumor necrosis factor-α (TNF-α) and interferon-γ (IFN-γ) mRNA levels increasing 3.0-fold (p<0.001) and 6.2-fold (p<0.05), respectively, compared to controls (Figure 3G-H). These findings collectively demonstrate prolonged intestinal susceptibility to *C.R.* infection through 7 days post-PEG exposure.

Notably, animals challenged at 14 days post-PEG administration showed complete resolution of infection susceptibility, with no significant differences observed in fecal bacterial burden (Figure 3J), colonic histopathology scores (Figure 3K-L), or cytokine expression profiles (Figure 3M-N) compared to untreated controls. This temporal resolution of pathogen susceptibility suggests full restoration of intestinal barrier homeostasis and microbial resistance mechanisms by 14 days following PEG-induced bowel preparation.

### PEG-induced alterations in gut microbiota composition

Given that intestinal cleansing has been shown to perturb gut microbiota, we conducted comparative analysis of fecal samples collected from mice at baseline (pre-PEG) and 1-day post-PEG using 16S rRNA gene sequencing. Principal coordinate analysis (PCoA) at the OTU level revealed distinct clustering patterns between pre- and post-PEG groups (Figure 4A). ANOSIM analysis confirmed significant intergroup differences in microbial community structure (R = 1.0, p = 0.004). Alpha diversity metrics demonstrated substantial reduction following PEG administration, with significant decreases observed in Chao index(p<0.001), Shannon index(p<0.001), and Ace index (p<0.001) at the OTU level (Figure 4B-D).

**Figure 4.**
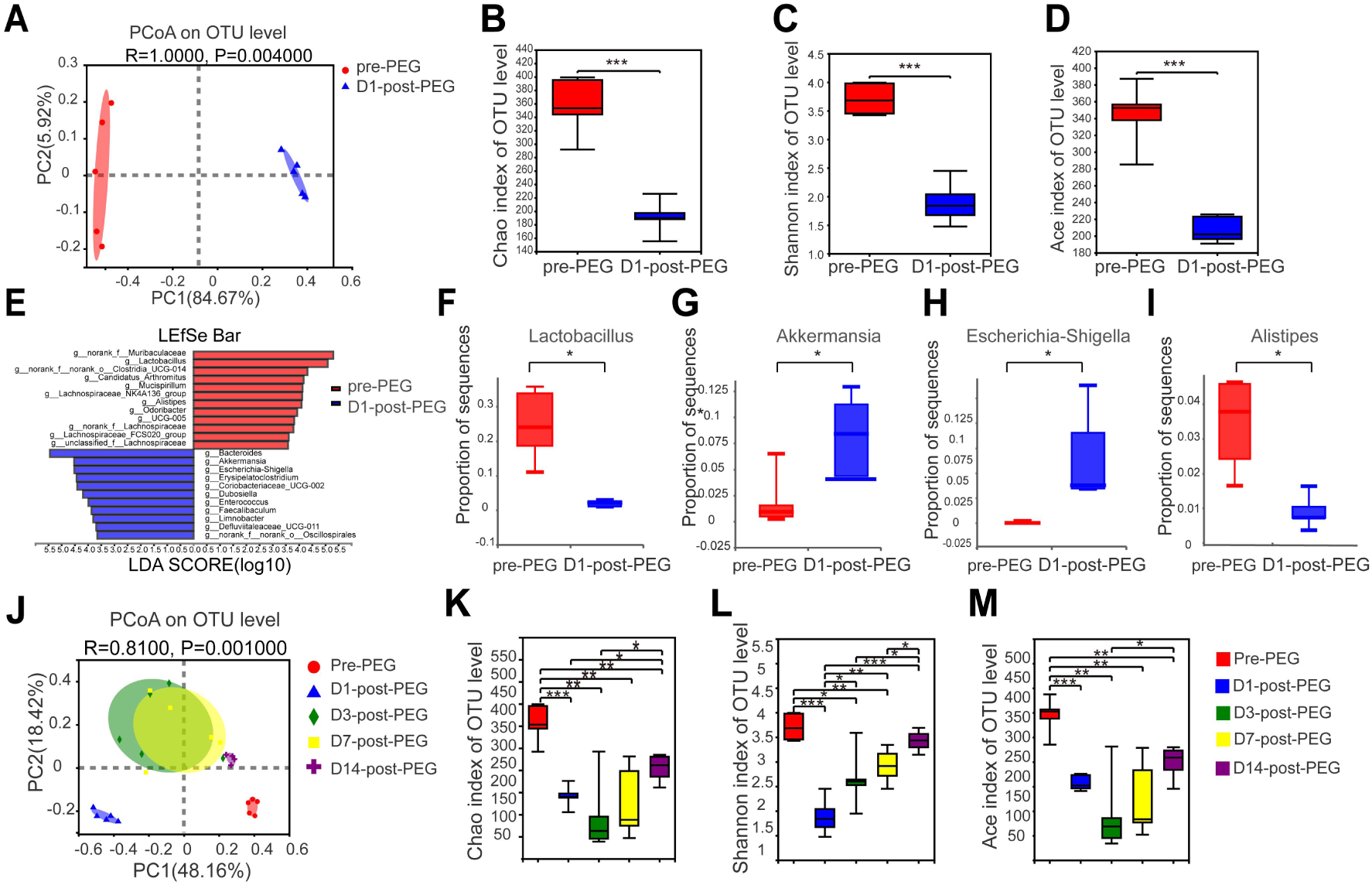
Gut microbiota dysbiosis play an essential role in the PEG relative infection. (A) Principal coordinate analysis (PCoA) of fecal microbiota pre-PEG and 1 day post PEG. (B-D) Shannon, Chao, and Ace indices of operational taxonomic unit (OTU) levels of fecal microbiota. (E) LDA score for differences between pre-PEG and 1 day post PEG. (F-I) Relative abundance of *Lactobacillus*, *Akkermansia*, *Escherichia-Shigella* and *Alistipes* on genus levels. (J) PCoA of fecal microbiota pre-PEG, 1 day post PEG, 3-day post PEG, 7-day post PEG and 14-day post PEG. (K-M) Shannon, Chao, and Ace indices of OUT levels of fecal microbiota pre-PEG, 1 day post PEG, 3-day post PEG, 7-day post PEG and 14-day post PEG. Data are expressed as mean ± SD; statistical significance was determined by a two-sided Student’s t-test. \**p* < 0.05; \*\**p* < 0.001; \*\*\**p* < 0.001. ns, no significance.

LEfSe analysis identified differential taxonomic enrichment between groups: Pre-PEG microbiota showed significant representation of *Muribaculaceae*, *Clostridia*, *Mucispirillum*, *Candidatus Arthromitus*, *Lachnospiraceae*, and *Alistipes*, while post-PEG communities exhibited predominance of *Bacteroides*, *Akkermansia*, *Escherichia-Shigella*, *Erysipelatoclostridium*, *Coriobacteriaceae*, *Enterococcus*, *Faecalibaculum*, and *Limnobacter* (Figure 4E-I).

To assess longitudinal recovery of microbial communities, fecal samples were collected at pre-PEG, 1-, 3-, 7-, and 14-day post-PEG intervals. Both β-diversity and α-diversity metrics demonstrated gradual temporal restoration, indicating partial microbiota recovery by day 14 (Figure 4J-M).

### PEG enhances colonic goblet cell proliferation

The macromolecular mucin MUC2, predominantly secreted by goblet cells, constitutes the colonic mucus layer. Histological evaluation revealed significant increases in goblet cell density following PEG preparation compared to baseline (p<0.001; Figure S1). This cellular proliferation suggests potential activation of negative feedback mechanisms to accelerate mucin production and restore mucosal barrier integrity.

### PEG-mediated susceptibility to C.R. infection is microbiota-dependent

To investigate whether PEG-induced gut microbiota alterations mediate increased susceptibility to *C.R.* infection, we performed a co-housing experiment where PEG-treated and untreated mice were housed in a 1:1 ratio for 7 days prior to *C.R.* challenge (Figure 5A). Notably, co-housed mice showed no significant differences in fecal *C.R.* colonization (Figure 5B) or colonic histopathological scores (Figure 5C-D) between PEG-treated and control groups.

**Figure 5.**
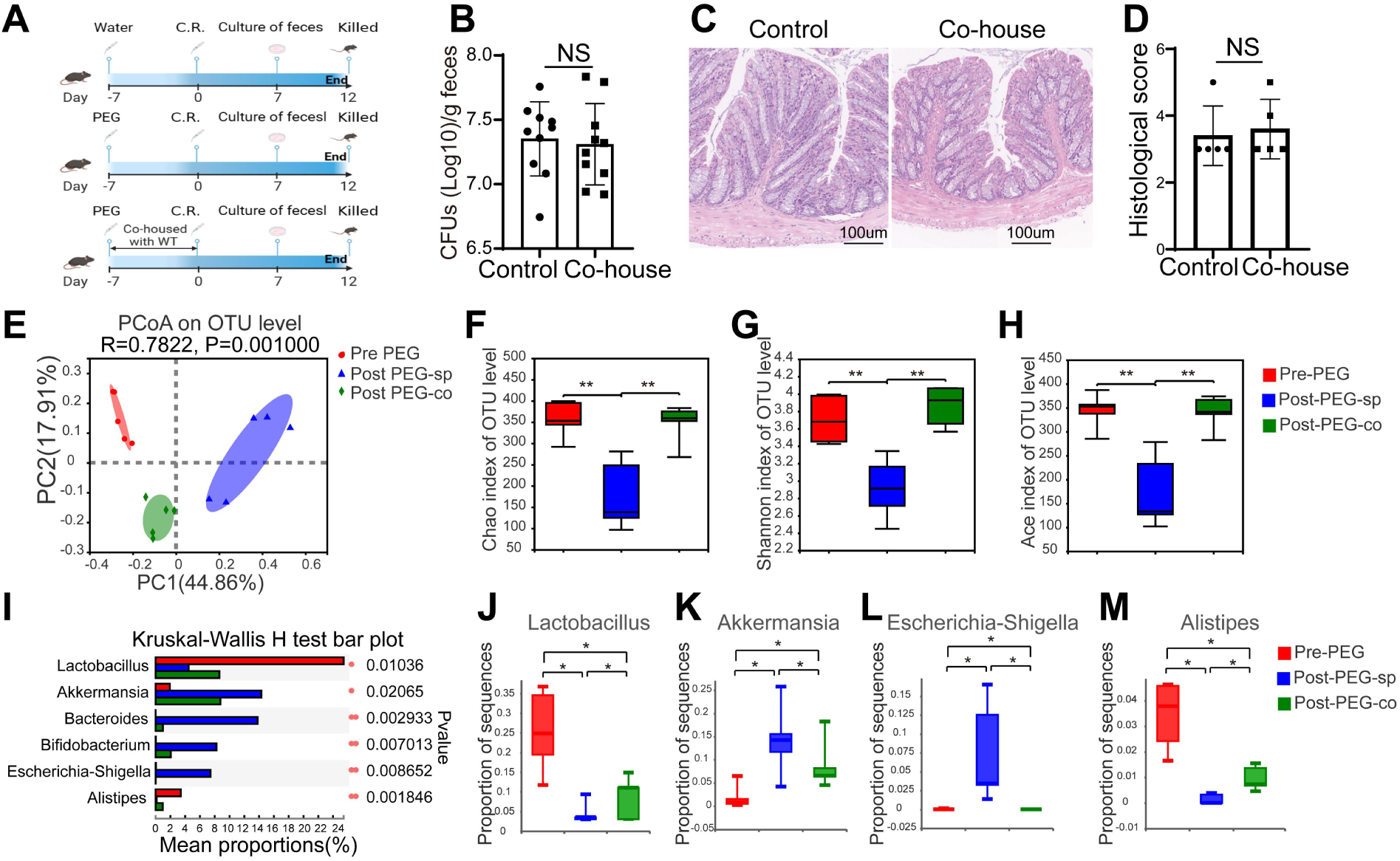
Co-house for 7 days could eliminate the effect of PEG. (A) Mice were separately housed and co-housed for 7 days post PEG administration and prior to the infection of C.R.. (B) C.R. load in the feces. (C-D) Representative H&E staining (×20) and pathological score of the colon epithelium. (E) PCoA of fecal microbiota among pre-PEG, 7-day post PEG of separately house and 7-day post PEG of co-house. (F-H) Shannon, Chao, and Ace indices of OTU levels of fecal microbiota among 3 groups. (I) Kruskal–Wallis H test for differences among 3 groups. (J-M) Relative abundance of Lactobacillus, Akkermansia, Escherichia-Shigella and Alistipes on genus levels. Datas are expressed as mean ± SD; statistical significance was determined by a two-sided Student’s t-test. *p < 0.05; **p < 0.01; ***p < 0.001. ns, no significance.

16S rRNA sequencing analysis of microbial communities revealed distinct clustering among three experimental groups: pre-PEG baseline, post-PEG separately housed, and post-PEG co-housed mice (Figure 5E). Co-housed mice exhibited microbiota profiles that converged toward pre-PEG baseline states compared to separately housed counterparts. Alpha diversity metrics including Shannon index, Ace index, and Chao index demonstrated complete restoration of microbial richness in co-housed mice to pre-PEG levels (Figure 5F-H).

Taxonomic analysis identified significant microbiota remodeling following co-housing, characterized by increased relative abundance of *Lactobacillales* (p<0.01) and *Alistipes* (p<0.05), coupled with marked reduction in *Escherichia-Shigella* populations (p<0.001; Figure 5I-M).

### *Lactobacillus* supplementation attenuates PEG-enhanced *C.R.* colonization

Given the observed depletion of *Lactobacillus* following PEG preparation, we investigated prophylactic administration of *LAC* on *C.R.* susceptibility. Mice received 1.86×10⁶ CFU/day*LAC* or PBS control for 3 days prior to PEG treatment (Figure 6A). *LAC* pretreatment significantly reduced fecal *C.R.* burdens compared to controls (p<0.001; Figure 6B) and ameliorated colonic inflammation as evidenced by lower histopathological scores (p<0.05; Figure 6C-D).

Microbiota analysis demonstrated distinct cluster separation among groups, with *LAC*-treated mice exhibiting closer to pre-PEG baseline than PEG-only controls (Figure 6E). In addition, *LAC* administration significantly enhanced microbial diversity as measured by Shannon index (p<0.001). Together, *LAC* application regulates the gut microbiota and mitigates PEG-increased *C.R.* infections.

**Figure 6.**
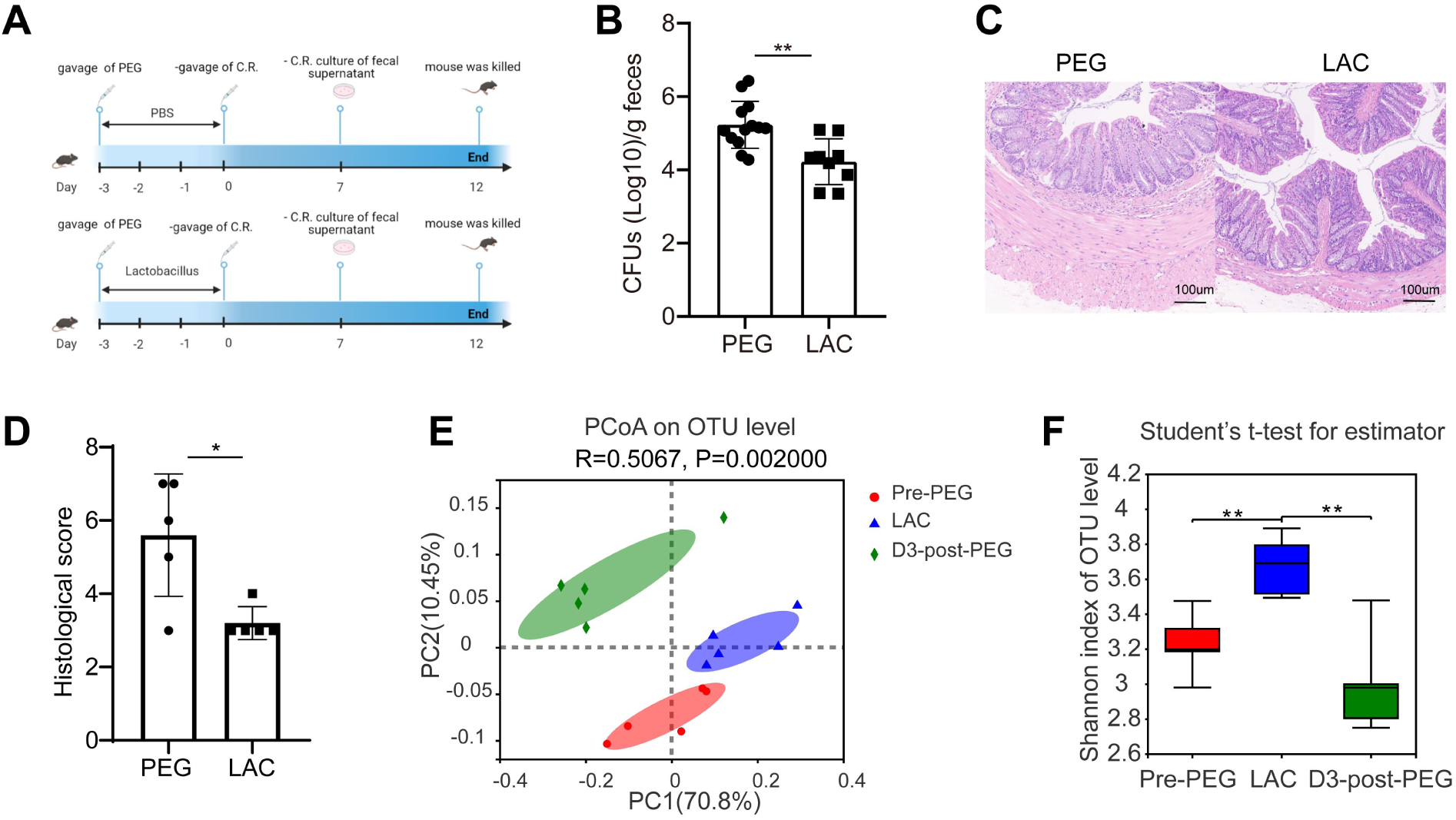
Supplementation of *Lactobacillus acidophilus (LAC)* could decrease the effect of PEG. (A) Mice were gavaged with PEG and treated with PBS or *LAC* for 3 days prior to the infection of *C.R*.. (B) *C.R.* load in the feces. (C-D) Representative H&E staining (×20) and pathological score of the colon epithelium. (E) PCoA of fecal microbiota among pre-PEG, 3-day post PEG and 3-day treatment of *LAC*. (F) Shannon indices of OTU levels of fecal microbiota among pre-PEG, 3-day post PEG and 3-day treatment of *LAC*. Dataare expressed as mean ± SD; statistical significance was determined by a two-sided Student’s t-test. \**p* < 0.05; \*\**p* < 0.01.

## Discussion

Adequate bowel cleansing remains essential for both diagnostic colonoscopy efficacy and surgical outcomes. While PEG-based solutions are widely regarded as safe and effective for this purpose [1, 16], their potential impact on enteric pathogen susceptibility has not been extensively investigated. This study demonstrates that PEG-induced bowel preparation elevates intestinal infection risk through by altering the intestinal microbiota, as well as impairing intestinal barriers. Notably, prophylactic administration of *LAC* mitigated these effects by restoring microbial homeostasis and reducing inflammatory responses.

Previous report suggested low-dose PEG (0.4 g/kg) may inhibit *C.R.* adhesion to colonic epithelium, highlighting its therapeutic potential against enteropathogens like *EPEC* and *EHEC* [17]. In contrast, our findings reveal that high-dose PEG administration paradoxically enhances bacterial colonization and virulence expression. Mechanistically, rapid electrolyte depletion and osmotic imbalance caused by PEG-induced diarrhea likely disrupt intestinal homeostasis, creating a permissive microenvironment for pathogen proliferation [18]. PEG consumption, washing out the luminal content, led to disturbances of intestinal fluid and osmolality. The mucus layer is an important component of the intestinal barrier function.

Defects of the mucus layer allow bacteria to reach epithelial cells more often than normal, thus activating the immune system (3). It has been established that the application of large amounts of PEG leads to thinning of the mucus layer and requires a longer time for recovery (4). Collectively, the substantial depletion of intestinal fluids and the concomitant reduction in mucus layer integrity were identified as critical determinants of enhanced infection susceptibility.

Consistent with established infection kinetics, both control and PEG-treated cohorts exhibited comparable fecal *C.R.* colonization patterns, reaching peak burdens at 7–8 d.p.i. followed by gradual clearance. This temporal similarity suggests PEG exacerbates infection severity without altering C.R. pathogenesis.

Our microbiota analysis revealed transient but significant PEG-induced dysbiosis, with full compositional restoration occurring by day 14 post-treatment. This 14-day "infection vulnerability window" correlates with dynamic fluctuations in mucin-associated taxa.

Specifically, we observed transient *Akkermansia* enrichment post-PEG-a genus positively associated with goblet cell density and barrier reinforcement in murine models [20–22]. This compensatory response may represent a host-protective mechanism counteracting PEG-induced mucosal damage. Published studies have demonstrated a significant reduction in *Akkermansia* abundance following a 6-day low-dose PEG administration protocol, with subsequent recovery observed upon treatment cessation. In our current investigation, we notably identified a marked elevation in *Akkermansia* relative abundance within the gut microbiota 24 hours post-PEG intervention. This microbial shift exhibited a significant positive correlation with increased goblet cell proliferation, suggesting a potential protective mechanism employed by the host organism to maintain intestinal mucosal homeostasis during osmotic challenge.

The therapeutic potential of *LAC* supplementation merits particular emphasis. As a well-characterized probiotic, *Lactobacillus* spp. enhance epithelial integrity through multiple mechanisms: tight junction modulation, antimicrobial peptide induction, and Wnt/β-catenin-mediated epithelial regeneration [23–25]. Our findings align with these mechanisms, demonstrating that *LAC* pretreatment reverses PEG-induced *Lactobacillus* depletion, reduces *C.R.* colonization, and attenuates colitis severity.

While murine models provide mechanistic insights, clinical translation requires validation in human cohorts. We acknowledge study limitations including the absence of large-scale clinical correlation data - a critical next step given the widespread use of PEG in clinical practice. Nevertheless, our results propose a pragmatic preventive strategy: short-term probiotic supplementation during bowel preparation protocols may reduce infection risks in vulnerable populations.

In summary, this study establishes that PEG bowel preparation transiently increases intestinal infection susceptibility through microbiota disruption and barrier impairment. Key findings include: PEG induces a 14-day vulnerability window characterized by dysbiosis and mucosal thinning. *Lactobacillus* depletion mediates enhanced *C.R.* colonization post-PEG. Prophylactic *LAC* administration restores microbial balance and reduces pathogen burden.

## List of abbreviations

PEG: Polyethylene glycol
*C.R.*: *Citrobacter rodentium*
*LAC*: *Lactobacillus acidophilus*
*EPEC*: *Enteropathogenic Escherichia coli*
*EHEC*: *Enterohemorrhagic Escherichia coli*
PPI: Proton pump inhibitors
CFUs: Colony forming units
MLNs: Mesenteric lymph nodes
H&E: Hematoxylin and eosin
PCoA: Principal coordinate analysis
OUT: Operational taxonomic unit
PBS: Phosphate buffer saline
D.p.i.: day post infection

## Declarations

### Ethics approval and consent to participate

The study received ethical approval from the Institutional Animal Care and Use Committee of Qilu Hospital, Shandong University (Approval No. Dull-2021-058). All procedures and methods adhered to the relevant guidelines and regulations.

### Consent for publication

Not applicable.

### Availability of data and materials

All data generated or analyzed during this study are included in this published article. The datasets generated of the study can be found in the National Center for Biotechnology Information (NCBI) database with accession code PRJNA1130659.

### Authors’ contributions

GK designed the study, analyzed and interpreted the data, and wrote the draft manuscript. JS, SF and LL participated in data collection and laboratory examinations. XZ and YL participated in data analysis and critically reviewed and drafted the final manuscript. All authors read and approved the final manuscript.

## Acknowledgements

We thank Prof. Shiyang Li, Mengqi Zheng and Zixiao Zhao for providing guidance and antibodies.

## Supplementary Materials

Figure S1: PEG increases the proliferation of goblet cells in colon.

