## Supplemental Figure 1 for "*Lactobacillus acidophilus* Attenuates Polyethylene Glycol-Induced Susceptibility to *Citrobacter rodentium* Infection via Microbiota Modulation"

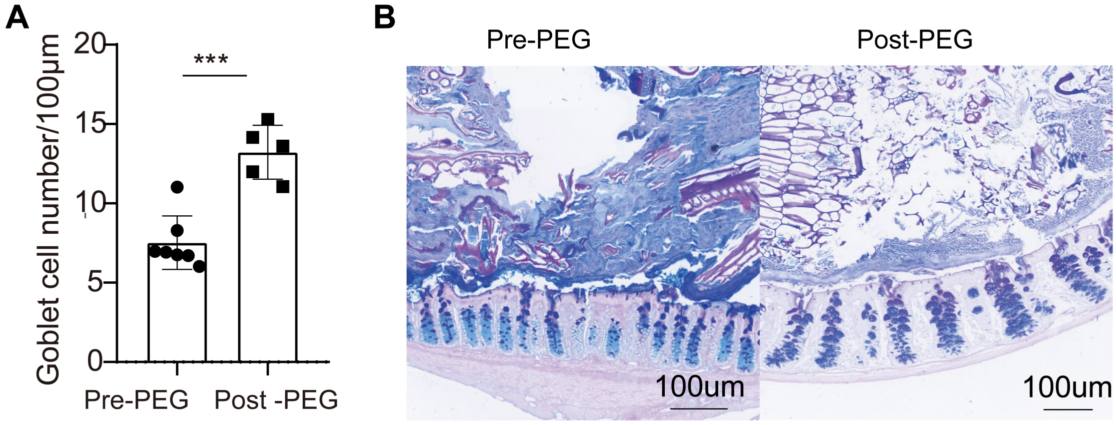


Supplemental Fig. 1 PEG increases the proferation of goblet cells in colon.

(**A-B**) PEG increases the proliferation of goblet cells in colon. Alcian blue staining of additional sections of the colon resulted in the blue staining of the cytoplasm of the goblet cells lining the intestinal epithelium, goblet cells were determined by Alcian staining colon tissue. Datas are expressed as mean ± SD; statistical significance was determined by a two-sided Student's t-test. ***$p < 0.001$.
